# An information theoretic treatment of sequence-to-expression modeling

**DOI:** 10.1101/316752

**Authors:** Farzaneh Khajouei, Saurabh Sinha

## Abstract

Studying a gene’s regulatory mechanisms is a tedious process that involves identification of candidate regulators by transcription factor (TF) knockout or over-expression experiments, delineation of enhancers by reporter assays, and demonstration of direct TF influence by site mutagenesis, among other approaches. Such experiments are often chosen based on the biologist’s intuition, from several testable hypotheses. We pursue the goal of making this process systematic by using ideas from information theory to reason about experiments in gene regulation, in the hope of ultimately enabling rigorous experiment design strategies. For this, we make use of a state-of-the-art mathematical model of gene expression, which provides a way to formalize our current knowledge of cis- as well as trans-regulatory mechanisms of a gene. Ambiguities in such knowledge can be expressed as uncertainties in the model, which we capture formally by building an ensemble of plausible models that fit the existing data and defining a probability distribution over the ensemble. We then characterize the impact of a new experiment on our understanding of the gene’s regulation based on how the ensemble of plausible models and its probability distribution changes when challenged with results from that experiment. This allows us to assess the ‘value’ of the experiment retroactively as the reduction in entropy of the distribution (information gain) resulting from the experiment’s results. We fully formalize this novel approach to reasoning about gene regulation experiments and use it to evaluate a variety of perturbation experiments on two developmental genes of *D. melanogaster*. We also provide objective and ‘biologist-friendly’ descriptions of the information gained from each such experiment. The rigorously defined information theoretic approaches presented here can be used in the future to formulate systematic strategies for experiment design pertaining to studies of gene regulatory mechanisms.

**Author summary:** In-depth studies of gene regulatory mechanisms employ a variety of experimental approaches such as identifying a gene’s enhancer(s) and testing its variants through reporter assays, followed by transcription factor mis-expression or knockouts, site mutagenesis, etc. The biologist is often faced with the challenging problem of selecting the ideal next experiment to perform so that its results provide novel mechanistic insights, and has to rely on their intuition about what is currently known on the topic and which experiments may add to that knowledge. We seek to make this intuition-based process more systematic, by borrowing ideas from the mature statistical field of experiment design. Towards this goal, we use the language of mathematical models to formally describe what is known about a gene’s regulatory mechanisms, and how an experiment’s results enhance that knowledge. We use information theoretic ideas to assign a ‘value’ to an experiment as well as explain objectively what is learned from that experiment. We demonstrate use of this novel approach on two extensively studied developmental genes in fruitfly. We expect our work to lead to systematic strategies for selecting the most informative experiments in a study of gene regulation.

## Introduction

Cellular processes are determined by the response of regulatory sequences in DNA to signals from specific proteins called transcription factors (TFs), leading to up- or down-regulation of gene expression [1]. A major class of regulatory sequences is that of cis-regulatory modules (CRMs, also called enhancers): these are regions of DNA, about 500-2000 base pairs long, harboring TF binding sites that control the transcriptional levels of nearby genes. Variation of the DNA sequence in CRMs can affect gene expression, and has been linked to developmental defects and disease [2]. Even minor variations, such as single nucleotide polymorphisms (SNPs), in CRMs can have significant functional impact, such as problems in fetal development [3].

Our ability to predict the impact of non-coding sequence variations on gene expression is very limited, in part due to the complexity of CRMs, and in part because such impact depends not only on the sequence itself but also the abundance and activities of relevant TFs in the cellular conditions of interest. Statistical methods based on correlations among diverse data types such as TF-ChIP, histone modifications, gene expression, etc. can reveal salient properties of CRMs such as their tissue-specific activities [4] and their major regulators [TFs] [5,6], and can in some cases predict the effect of removing a TF’s influence on a CRM or gene [7–9]. Statistical and machine learning methods have recently been developed that can to some extent predict the effects of single nucleotide mutations on TF binding levels, DNA accessibility [10,11], and even gene expression [12], but these are typically not amenable to mechanistic interpretations, and are in a relatively early stage of exploration.

On the other hand, biophysical models based on equilibrium thermodynamics that explicitly incorporate key interactions among TFs, DNA and the transcriptional machinery have proven powerful for mechanistic understanding of the gene regulation process [13]. Thermodynamics-based modeling of gene expression reveals the precise mapping between CRM sequence and the associated gene expression in a variety of cellular contexts, the so called ‘readout’ of the CRM. These models provide a means to formalize our assumptions about a CRM’s cis-regulatory logic, especially how its functional elements combine to regulate a transcriptional output [14–19]. They can generate predictions that can be empirically tested [20], e.g., by targeted misexpression or mutagenesis experiments. Indeed, they have been used to predict effects of site mutations [21], and also promise to provide precise, mechanistically grounded predictions of the effect of minor sequence changes in CRMs [20]. Furthermore, these models can reveal ambiguities in our mechanistic knowledge about a system given existing data; pinpointing these ambiguities helps with choosing the future experiments that would best improve knowledge of the system. The success of thermodynamic models has been demonstrated in the context of systems with high-resolution gene expression measurements, such as early-stage Drosophila embryonic development [15–17, 22].

Mechanisms influencing the precise function of a regulatory system include the number, accessibility, affinities and relative arrangement of TF binding sites within a CRM, as well as cellular concentrations of the TF molecules, and protein-protein interactions; all of these mechanisms affect the rate of transcription of the gene [23]. Thermodynamic models of CRM function encode these mechanistic factors in their parameters, which correspond to biochemical properties of the molecules controlling the gene expression. These parameters are typically computationally optimized to be assigned values that can best explain the gene expression patterns attributable to a set of CRMs [16,22]. When used to investigate the regulatory function of a single CRM, the thermodynamic modeling approach faces a significant challenge: non-uniqueness of the optimal models. For instance, a CRM mediating control by five or more TFs will include 10 or more free parameters in the GEMSTAT thermodynamic model [22] and we have shown previously that parameter training will converge to one of many local optima [21]. Each optimal model explains the data equally well, but uses parameters that correspond to significantly varying, often mutually incompatible mechanistic hypotheses [24,25]. For example, consider a gene regulated by a CRM that is under the control of two activators. Assume there are two models that explain the wild-type expression pattern. One model predicts the correct expression by using (assigning function to) only one activator, while the other model uses both activators. In the absence of additional biological experiments that confirm the role of each activator, both models are equally plausible. Problems arise when we try to predict, using the model, the effect of knocking down an activator or mutating its binding site(s). Depending on which model we use, the predicted effect of the perturbation experiment is different: a model that does not use an activator will not predict a change due to removal of that activator’s influence. However, there is no reason to prefer one model’s prediction over the other, and the biology remains ambiguous until a new experiment is performed. We believe it is important to respect this ambiguity of knowledge when modeling gene expression data and making predictions about future experiments.

In agreement with the above proposal, Samee et al. [21] laid out a new paradigm of gene expression modeling where one searches within the model’s parameter space for as many optima as possible, resulting in an “ensemble” of optimal models. (Henceforth, different assignment of values to the model’s tunable parameters will be considered as different models.) Each model in the ensemble is a hypothesis about the cis-regulatory mechanisms encoded in the CRM, and is also capable of making specific predictions about perturbation experiments. A simple approach to working with an ensemble of models is to make predictions by uniformly aggregating predictions of its member models. It has been shown that this “wisdom of crowds” approach can be effective: aggregated votes of many models can predict the effect of site mutations more accurately than any individual model [21].

We noticed, however, that a typical ensemble of sequence-to-expression models, e.g., that created by Samee et al. [21] in modeling embryonic expression of the *Drosophila* gene *ind*, is not uniformly distributed in the parameter space. Rather, they are clustered in the parameter space (Fig 1D), with models within a cluster predicting similar effects for a particular perturbation, but each cluster’s consensus predictions being qualitatively different from those of other clusters. Different clusters can have different ‘spans’, i.e., the extent to which models in that cluster differ from each other quantitatively (in parameter values) while producing equally good fits to available data and essentially the same predictions for future experiments. For instance, the cluster at the bottom of Fig 1D has greater span than the cluster on the top-left, which is relatively tight. The span of a cluster pertains to parameter sensitivity [24,26] in that region of parameter space.

**Fig 1.**
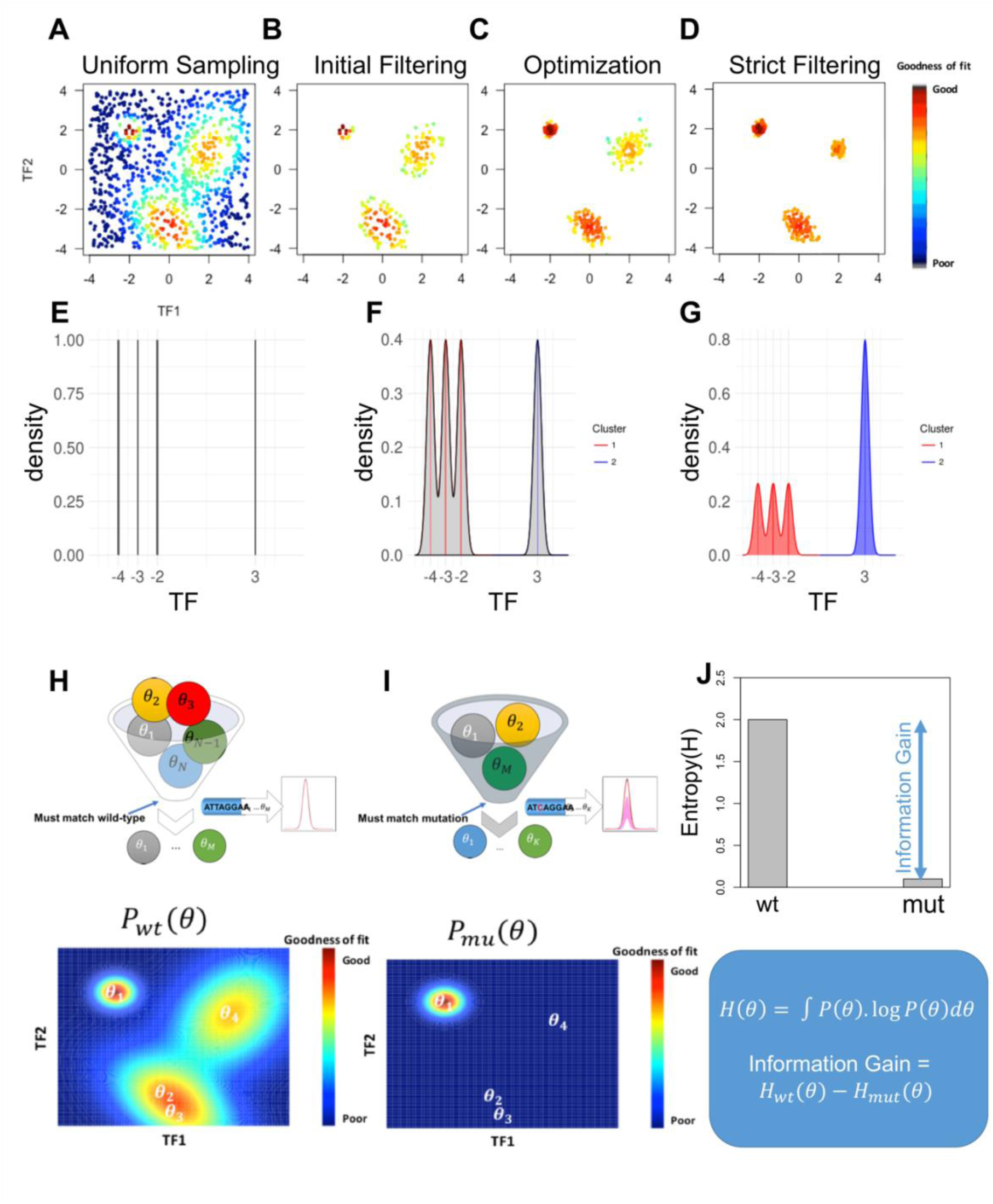
Schematic of Methodology. **(A)** Each point in the scatter plot is a model with two parameters denoted TF1 and TF2 and its color shows its ‘goodness-of-fit’ score (see color legend on right). A number of models are uniformly sampled from the two-dimensional parameter space and scored against available data. **(B)** Initial filtering of models in (A) for a high goodness-of-fit score results in multiple clusters of models. **(C)** Models in panel B are used as starting points for numeric optimizations of goodness-of-fit, resulting in multiple clusters of locally optimal models. **(D)** A stringent threshold on goodness-of-fit is used to filter optimized models (panel C) to obtain the final ensemble. **(E)** An ensemble of four models in a one-dimensional space is represented by a uniform discrete probability distribution. **(F)** The same ensemble as (E), represented by a continuous probability distribution, as a uniform mixture of Gaussian distributions centered at the four models. **(G)** Models are recognized to fall in two clusters and the continuous distribution of (F) is modified to assign equal weight to each cluster. **(H-I)** Illustration of an information theoretic value of an experiment: (H) All models are assessed for goodness-of-fit on available (e.g., wild-type) data, an ensemble (θ_1_, θ_2_,… θ_M_) is obtained (top) as outlined in panels A-D, and a probability distribution is constructed (bottom), shown here for a two-dimensional parameter space. (I) Models in the wild-type ensemble are further examined for goodness-of-fit on new experimental data (e.g., a site mutagenesis experiment), and fit models are retained to construct a new filtered ensemble (top) and the probability distribution is recomputed (bottom). (J) Entropy scores of the probability distributions of the wild-type ensemble and filtered ensemble are computed, denoted by H*_wt_* and H_*mut*_ respectively, and the information gain is computed as the difference of these two entropy scores.

Furthermore, different clusters may have different representation (number of models) in the ensemble, and the number of represented models may not correlate with the span of the cluster. This is because we do not make strong assumptions about how the ensemble of models was obtained, beyond that it is a collection of models that fit the available data and may be located in different regions of parameter space. With these observations about ensembles of models, we sought the most appropriate way to use ensembles for making predictions and for designing future experiments. We describe here one such procedure that we developed and implemented, which allows us to make predictions with ensembles of models, and also offers a principled approach to experiment design in gene regulation studies.

Briefly speaking, our modeling approach involves (1) creating a large ensemble of models that fit the available data accurately, following the sampling and optimization strategy of Samee et al [21] and (2) defining a probability distribution over the parameter space such that the ensemble of models represents regions of high probability and where each cluster of models (roughly speaking, a distinct mechanistic hypothesis) has approximately the same total probability as other clusters. This distribution provides a principled way for us to make aggregated predictions about any particular perturbation experiment, and to describe the uncertainty in such predictions. Additionally, we show how to measure the entropy of this probability distribution, thereby quantifying the uncertainty in parameter space [27] that remains after fitting the models to available data. Noting that the ensemble of models consistent with available data changes (typically shrinks) upon performing an additional experiment, we suggest that the difference of entropies of the probability distributions before and after an experiment (i.e., information gain) may be used to score the ‘value’ of the experiment. We can use this value as a score to compare different experiments, the experiment with greater score being deemed the more informative experiment. The ability to assign information theoretically-grounded ‘values’ to experimental results is significant, since it paves the way for principled experiment design [28,29]

## Results

### Construction of a probability distribution over models, and potential applications Outline of gene expression model

We consider the class of mathematical models that predict the gene expression level driven by a cis-regulatory module (CRM) from the latter’s sequence, given prior knowledge of relevant transcription factors (TFs), their *in vitro* DNA-binding affinities (motifs), and their concentration levels in the cellular context of interest. Several such models have been investigated in the literature [15–17, 20], and we work with the GEMSTAT model [22], which we developed previously and which we are most familiar with. The GEMSTAT model has two free (tunable) parameters for each relevant TF, one corresponding to its binding strength for the consensus site and one corresponding to its potency as an activator or repressor. The model also has optional free parameters for any TF-TF cooperative interactions that the modeler may choose to include. Assigning values to these free parameters specifies a model completely, allowing it to predict gene expression in any cellular context where TF concentrations are known. Typically, optimization strategies are used to identify the parameter setting(s) that accurately predict gene expression driven by a CRM in multiple cellular contexts [30].

### Construction of model ensemble

In light of the observations made in Introduction, we sought to first construct an ensemble of models that are widely spread in parameter space, and thus represent different mechanistic explanations of data. A model is included if its goodness-of-fit score – sum of squared errors or ‘SSE’ between known and predicted expression levels in multiple cellular conditions – is below a threshold. We noted that the number of TFs in common modeling scenarios is less than 10 [15–17,20,22], and the number of free parameters in the range of 10-20. This led us to consider uniform sampling of the parameter space as the first step of ensemble construction. We followed the approach of Samee et al. [21] (Fig 1 A-D), performing extensive uniform sampling from the space (millions of samples), followed by filtering of promising models (SSE score below a modest threshold), local optimization seeded by these promising models, and a final round of filtering on the optimized models (SSE score below a strict threshold). (See Methods for details.) This procedure allows us to construct a large ensemble of models representing many or all optimal regions of the parameter space. We provide more details of ensemble size and composition later, in the context of specific gene models.

### Construction of probability distribution over models

An ensemble of models can be used to make predictions by aggregating (averaging) the predictions made by each member model. However, this approach ignores the fact that the ensemble construction (outlined above or by a similar method) likely results in some regions of parameter space being over-represented in the ensemble. Models belonging to the same region, i.e., proximal to each other in the parameter space, are presumed to represent qualitatively similar mechanisms of CRM function. Thus, the ensemble’s aggregate predictions may be biased towards one or a few mechanistic hypotheses. We therefore sought a more nuanced way to aggregate model predictions, by defining a probability distribution over parameter space that captures how the fit models are spread across different regions of the space but discounts for unequal representations of (number of models in) different regions. Such a probability distribution can then be used to make predictions about new experiments and also to score the uncertainty of mechanistic explanations offered by the ensemble. We also note that constructing this distribution has close ties to the kernel density estimation problem [31] but is different because the ensemble is not a collection of IID samples drawn from the desired population.

The simplest distribution to consider is a discrete uniform distribution over the models in the ensemble; e.g., Fig 1E shows such a distribution over an ensemble of four models in a toy 1-dimensional parameter space. In the continuous parameter space, highly proximal models are likely to have similar goodness-of-fit, therefore we smoothen the discrete distribution by centering a Gaussian distribution at each model in the ensemble and constructing a uniform mixture (Fig 1F). This mixture of Gaussian distributions provides a continuous distribution, but if one region of the space is over-sampled in the ensemble, the distribution puts undue weight in that region; e.g., the three closely-related models on the left in Fig 1F together carry about three times the probability mass as that around the isolated model on the right. In light of this observation, we first cluster models, each cluster roughly corresponding to a distinct mechanistic hypothesis, and define the overall probability distribution to be a mixture of distributions representing each cluster. Since we lack any additional knowledge to prefer one cluster over another, we assume uniform mixture weights for the clusters. The probability distribution representing each cluster, in turn, is a mixture of Gaussian distributions whose means are the models in that cluster. Thus, Fig 1G shows a mixture of two distributions (red and blue) representing the two clusters, with the red distribution in turn being a uniform mixture of Gaussians centered on the three models in that cluster. Fig 1H shows a similar construction, now for a 2D parameter space, beginning with the given filtered ensemble (θ_1_, θ_2_,… θ_M_), identifying three clusters and constructing the mixture probability distribution. (For more details, especially the construction of covariance matrices for these distributions, see Methods.)

### Potential Applications

The probability distribution over models, constructed as above, can be used in the following ways:

1. Aggregating ensemble predictions: Each model in the ensemble makes a prediction on unseen data, i.e., the gene expression level driven by the modeled enhancer(s) in a cellular context described by TF concentrations. The ensemble’s aggregate prediction can be computed by averaging predictions of all models weighted by their probabilities as specified by the distribution.
2. Quantifying uncertainty of ensemble predictions: The probability distribution makes it possible to quantify the variance in ensemble predictions on unseen data. This variance represents the uncertainty among ensemble models with respect to that data point.
3. Assigning objective ‘value’ to an experiment: We can utilize the probability distribution over an ensemble to measure the information theoretic value of an experiment. Given new data, i.e., results from a new experiment, we can filter models to obtain a smaller ensemble that agrees with the new data, henceforth called the ‘filtered ensemble’ for the experiment. Using the fact that the entropy of the probability distribution captures the uncertainty intrinsic to the ensemble, we can use the difference in entropy of the original ensemble and the filtered ensemble, also called the ‘information gain’, as an objective evaluation of how informative the experiments results are. (Details of entropy calculation for a given probability distribution are provided in Methods, but see Fig 1H,I for an illustration.) This approach ultimately allows, though we do not explore it here, principled experiment design where the experiment whose results are expected to result in the greatest information gain, is selected for follow-up.

#### An information theoretic measure of the ‘value’ of an experiment

Sequence-to-expression models enable us to propose mechanisms for gene expression regulation, that may then be confirmed by performing perturbation experiments such as TF knockout or site mutagenesis, followed by expression assays that inform us about how the gene expression changes in the perturbation condition. Some experiments result in greater gene expression changes than others, and it is natural to want to characterize ‘what was learned’ from each experiment, as well as quantify how informative that experiment was. Here, we demonstrate such an exercise in systematic experimentation in the gene regulation context, using the ensemble modeling framework described above.

Our first set of demonstrations are in the context of the regulatory mechanisms of an early development gene in *D. melanogaster* - the intermediate neuroblasts defective (*ind*) gene. We chose this gene because it is known to be regulated by a well-defined enhancer, and its major regulatory inputs are well characterized. The gene was characterized by Weiss et al. [32] and Stathopoulos and Levine [33], among others, and was the subject of systematic modeling by Samee et al. [21]. It is expressed in a lateral stripe along the dorso-ventral axis of the early embryo (S1A Fig, black curve), with activation from the TFs Dorsal (DL) and Zelda (ZLD), and repression by the TFs Snail (SNA), Ventral nervous system defective (VND) and Capicua (CIC) (S1A Fig). In addition to the wild-type expression pattern of this gene, its expression has been experimentally recorded under several perturbation conditions (S1 Table), surveyed by Samee et. al [21] and further discussed below.

Despite the knowledge of a fairly complete set of regulatory inputs, several ambiguities remain about the cis-regulatory logic of the *ind* enhancer. This is evident when we construct an ensemble of models that predict the known expression pattern of *ind* from its enhancer sequence along with TF concentration profiles along the D/V axis. S1B Fig shows that the ensemble’s mean prediction (magenta curve) for these wild-type conditions fits the wild-type expression profile accurately, and with little variation among different models (pink curves are models in the ensemble) but S1C Fig reveals that most of the 13 parameters of the model exhibit substantial variability, a point also illustrated by the marginal distributions of ten of the parameters (S1D-F Figs). The high degree of uncertainty is not surprising, given that data from only one experiment – the wild type condition – for a single enhancer was used to train the ensemble. It also means that results of various perturbation experiments may prove informative about this gene’s regulatory mechanisms, an avenue that we pursue next.

First, we worked with a ‘synthetic true model’ M_ST_ that allows us to predict results of various perturbation ‘experiments’ *in silico*. This synthetic true model M_ST_ was carefully chosen from among the ensemble of models consistent with wild-type data, described above. (See S3 and S4 Figs for details.) We used M_ST_ to individually predict the effects of (a) each TF’s knockout and (b) removing the strongest site of each TF in the enhancer, and treated these predicted gene expression patterns (Fig 2A, green curves) as the ‘true’ results of those hypothetical or ‘*in silico* perturbation experiments’. We used each of the 10 *in silico* experiments to construct a ‘filtered ensemble’ (average predictions shown in Fig 2A, magenta curves), computed its entropy score, and thus assigned an information theoretic ‘value’ to the experiment (Fig 2B). We noted that the magnitude of change in the expression profile resulting from a perturbation experiment does not necessarily reflect the value of the experiment. For instance, it is possible to obtain new information from a perturbation experiment where the expression pattern remains unchanged from wild-type, a case in point being the SNA knockout experiment (Fig 2A), with assigned value 1.66 – apparently many models consistent with wild-type data cannot explain this experiment and are removed in the filtered ensemble from its results. Conversely, an experiment with a more substantial expression profile change may not add anything to our knowledge of the regulatory mechanism. For instance, the DL knockout experiment shows peak *ind* expression diminishing by ~60% (Fig 2A) but is assigned a value of 0.32 (Fig 2B), among the lowest of the 10 experiments; this is because most models capable of explaining the wild-type *ind* pattern apparently use DL as activator, so knocking out DL does not provide much new information. The same is not true of the experiment where the strongest DL site is removed, an experiment with minor impact on expression (Fig 2A) but a relatively high assigned value of 1.34. This points out that even if the involvement of a TF is beyond doubt, there may be uncertainty regarding the strength of its regulatory input and the mediatory role of each of its binding sites. We noted (Fig 2B) the same trend -- that the value of strongest site mutagenesis is greater than that of TF knockout – for the other activator (ZLD). On the other hand, for perturbations involving repressors (SNA, VND, CIC) the value of the site mutagenesis experiment is less than that of TF knockout in all three cases. Also, for comparison, we show in Fig 2C the relative values of the 10 ‘experiments’ under a more simplistic scheme that evaluates each experiment by the reduction in entropy assuming a discrete uniform distribution on all models in an ensemble. We note that the two schemes largely agree with each other in this evaluation, though this may not be true in general, depending on how an ensemble of models is generated. Finally, we note that the observations above were made with a specific choice of the ‘synthetic true model’ M_ST,_ that furnished ‘experimental’ results, but the reported trends, e.g., large information gain from a perturbation experiment with little effect on expression, or little gain from an experiment with large effect, were unchanged when we repeated the entire exercise with a different choice of M_ST_ ( S5 and S3B Figs).

**Fig 2.**
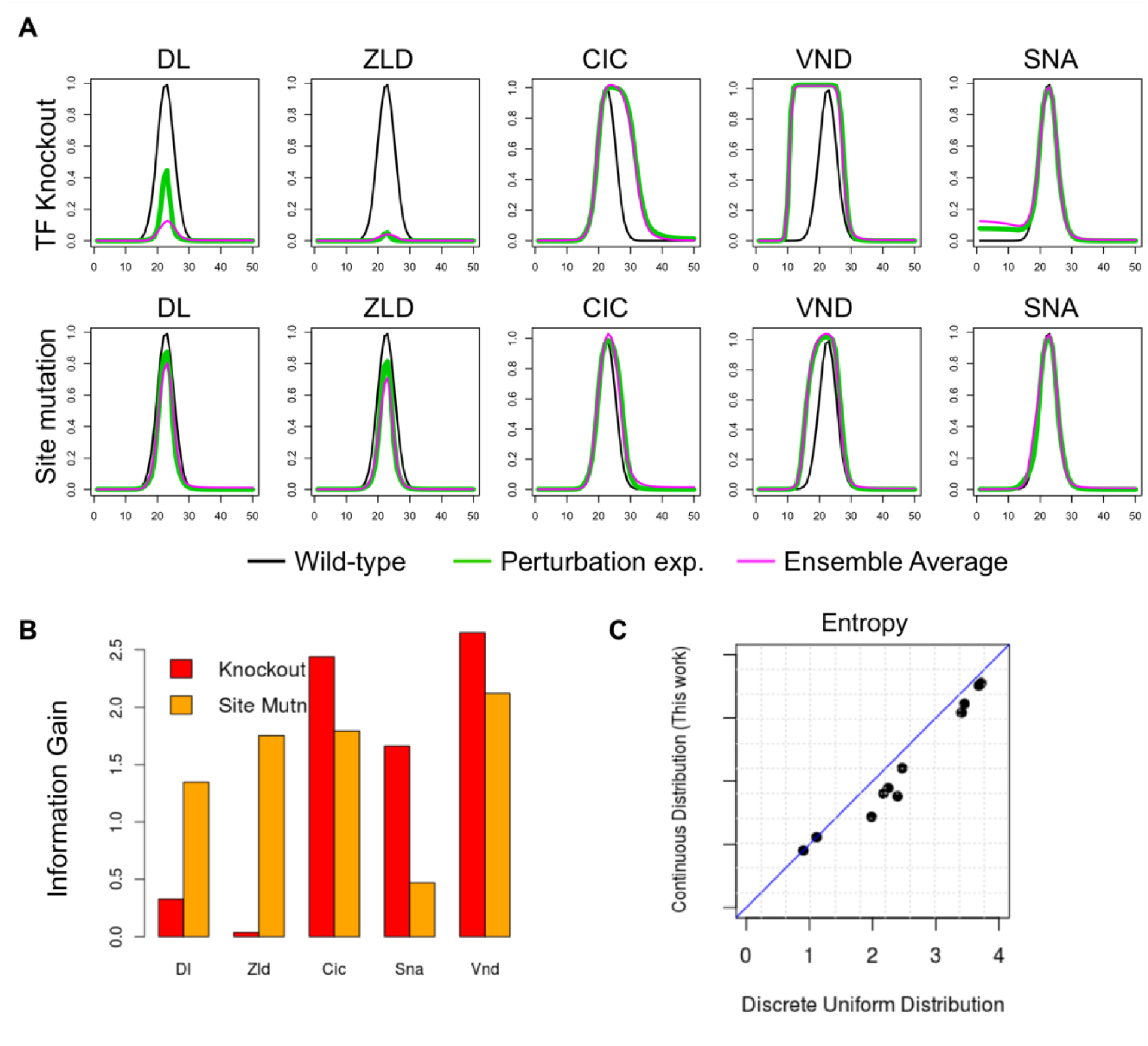
Evaluating in silico experiments with a ‘synthetic real’ model M_ST_. **(A)** The model is used to generate synthetic ‘experimental’ results of TF knockout (top row) or strongest site mutagenesis (bottom row), for each TF, shown in green. These are compared to the synthetic ‘wild-type’ expression profile of *ind*, shown in black (in each panel). Magenta curves show the average prediction of the filtered ensemble for each of these ‘experiments’. **(B)** Each of the ten synthetic ‘experiments’ (panel A) is assigned a value, which is defined as the information gain due to that experiment. **(C)** Entropy of filtered ensemble for each of the ten ‘experiments’, as defined under the specially constructed probability distribution presented in this work (Y axis) or under a discrete uniform distribution (X axis).

#### Evaluating perturbation experiments on *ind* gene regulation, *ex post facto*

In this section, we will examine results of real perturbation experiments pertaining to the *ind* gene reported in the literature and evaluate each experiment in the way described above. In addition to the wild type gene expression pattern of the *ind* gene (S1A Fig), we have information from six different biological perturbation experiments (S1 Table). It is known that *ind* expression is abolished in DL mutants [34] and becomes weaker in ZLD mutants [35]. Its peak expression reduces to ~50% of its wild-type level upon mutation of the four strongest ZLD binding sites [21]. (We call this experiment ‘ZLD site mut.’.) Removal of the strongest DL site (‘DL 1 site mut.’) has no observable effects on the expression [36] and removing three overlapping DL sites (‘DL 3 site mut.’) greatly diminishes peak expression [21]. Knockout of SNA (experiment ‘SNA KO’) leaves *ind* expression unaltered [17], while knocking out VND (‘VND KO’) causes the domain of expression to expand ventrally [37], and CIC site mutagenesis (‘CIC site mut.’) expands *ind* expression dorsally [38]. We evaluated each of the six perturbation experiments (two TF knockouts and four site mutagenesis experiments) using the approach introduced in the previous section – begin with the ensemble of models that explain wild-type gene expression, construct a filtered ensemble that additionally explains the perturbation results (see Methods and S2 Fig), and calculate the difference in entropy (‘information gain’). The values assigned to these experiments are shown in Fig 3A, and we note that the SNA and VND knockout experiments were the most informative in this group.

**Fig 3.**
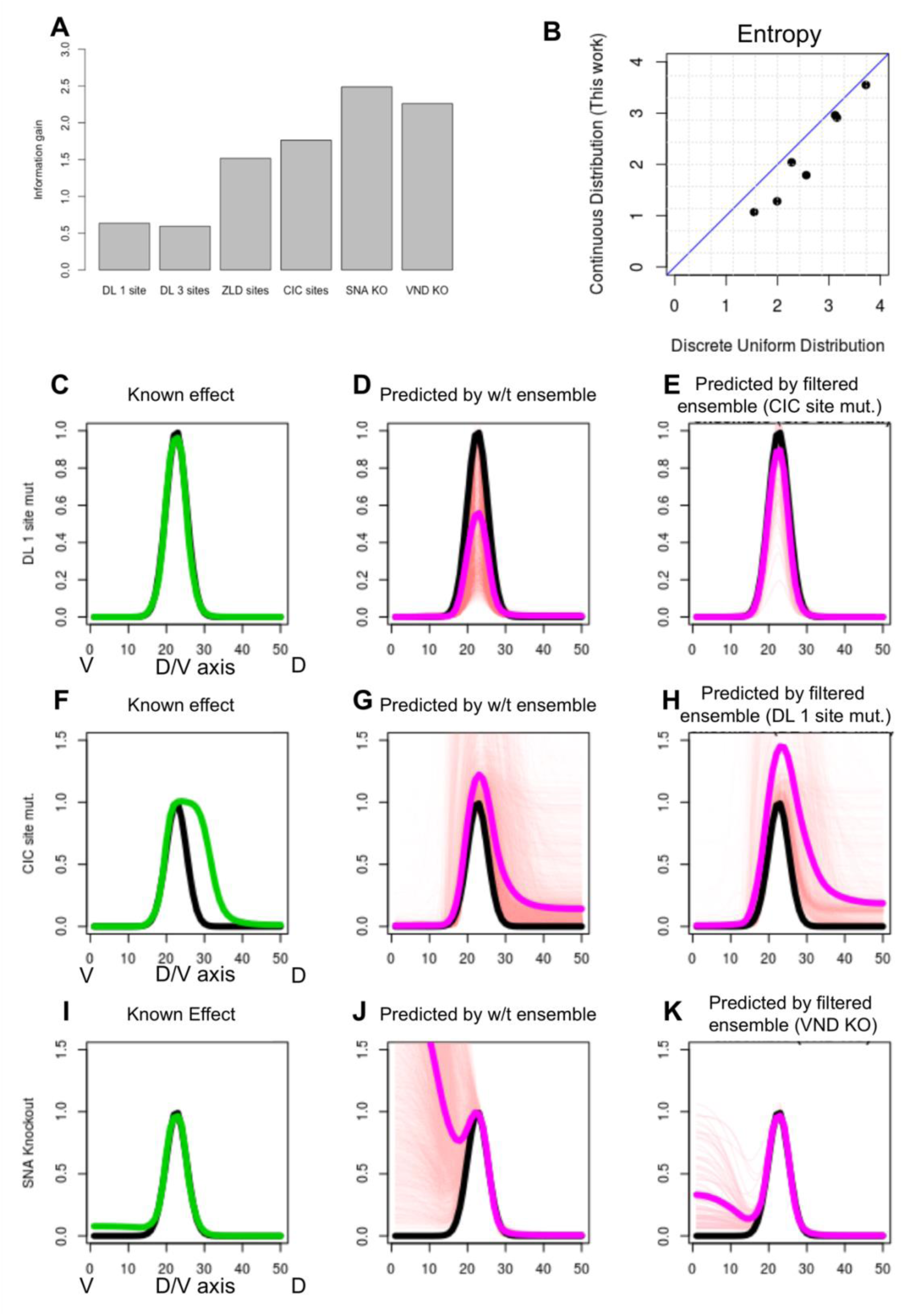
**(A)** Information gain score assigned to each of six perturbation experiments involving regulation of the *ind* gene. These include four site mutagenesis experiments (four left bars) and two TF knockout experiment (two right bars). **(B)** Entropy of filtered ensemble for each perturbation experiments, as defined under the specially constructed probability distribution presented in this work (Y axis) or under a discrete uniform distribution (X axis). **(C-E)** Known effect of the ‘DL 1 site mut.’ experiment (C, shown in green, compared to wild-type in black) is better predicted by the filtered ensemble for the experiment ‘CIC site mut.’ (E) than by the wild-type ensemble (D). In D, E, ensemble predictions shown as thin red lines and their mean shown in thick pink. **(F-H)** Known effect of the ‘CIC site mut.’ experiment (F) is better predicted by the filtered ensemble for the ‘DL 1 site mut.’ experiment (H) than by the wild-type ensemble (G), which shows greater variance and disagreement across models. **(I-K)** The ‘SNA KO’ experiment has been observed to not affect *ind* gene expression pattern (I). In contrast, majority of the models in the wild-type ensemble (J) predict substantial ventral de-repression, but an ensemble filtered by the ‘VND KO’ experiment predicts far less change in the ventral domain of *ind* expression profile.

Evaluating a new experiment, in our scheme, involves ruling out from the original ensemble a subset of models inconsistent with that new experiment. Recall that models in the ensemble were clustered, with the informal understanding that each cluster represents a distinct mechanistic hypothesis. Thus, if an entire cluster is ruled out by a particular experiment, one may interpret it as ruling out a particular mechanistic hypothesis. Table 1 shows the sizes of clusters in the original (wild-type) ensemble of models and the effect of filtering with each perturbation experiment. We note that an experiment (‘DL 3 site mut.’ in Table 1) may remove just one cluster, while retaining other clusters of models as feasible. There may also be experiments (‘SNA KO’ and ‘VND KO’ in Table 1) that rule out the majority of mechanistic hypotheses, retaining only 2-3 of the original clusters. The other scenario – where all clusters are retained but rendered substantially sparser – is also seen, indicating that the information gained by those experiments was more along the lines of quantitative refinement rather than qualitative pruning of the space of possible mechanisms. Fig 3B shows the above information theoretic evaluation of each experiment, compared to a simpler scoring scheme where entropy of an ensemble is simply the logarithm of the size of that ensemble, i.e., where we assume a uniform discrete distribution on models. As expected, the two scores are highly correlated.

**Table 1.**
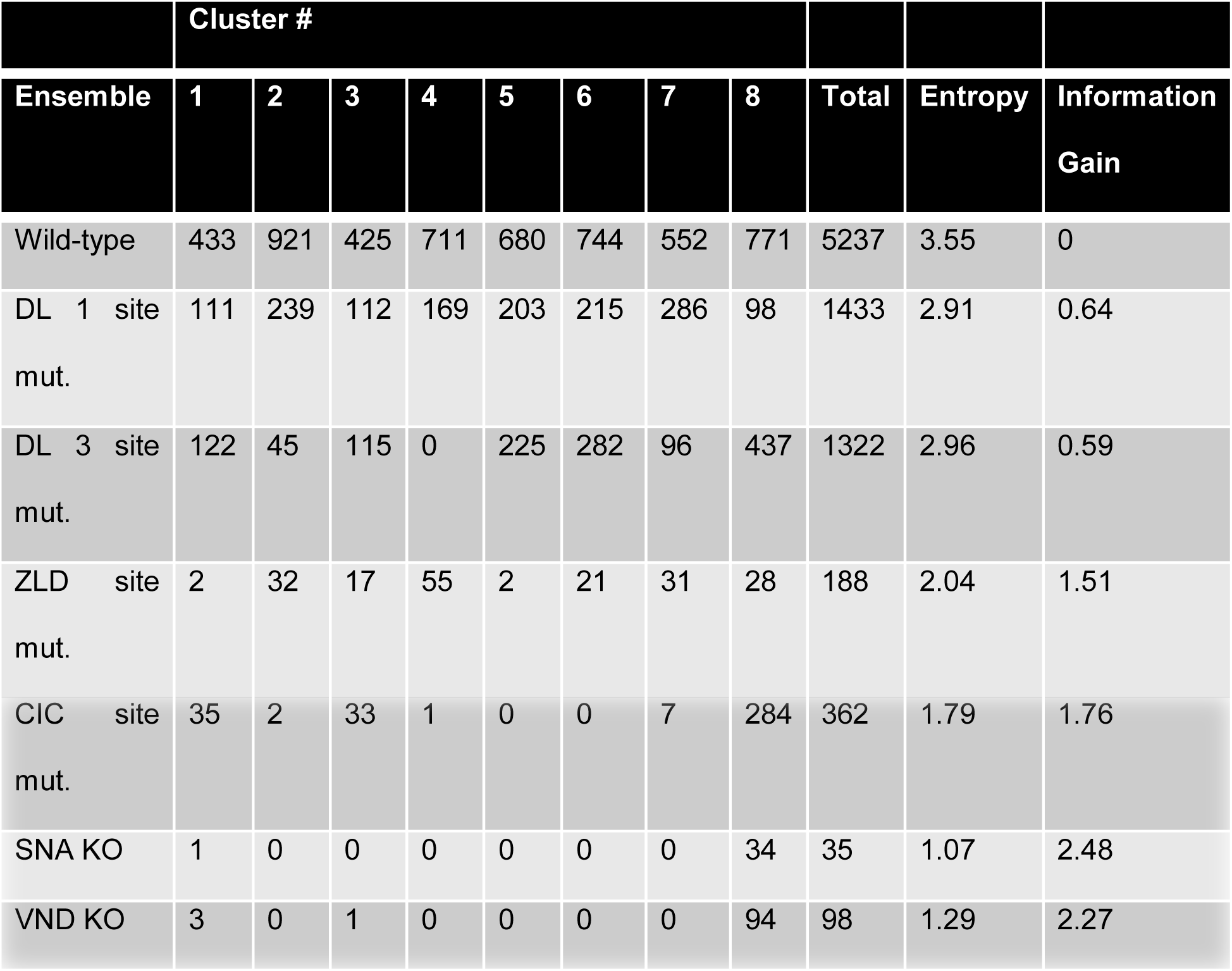
Models in the wild-type ensemble can be clustered into 8 different groups based on their parameter values, each cluster roughly corresponding to a distinct mechanistic hypothesis. Given information from a new experiment, we filter the wild-type ensemble for models that are predictive of the experiment outcome. Shown here are the sizes of clusters of models in the wild-type ensemble, and their corresponding sizes after filtering for each of six different perturbation experiments. The total number of models in each ensemble, the corresponding entropy score and information gain score are shown in the last three columns.

Note that experiments were assigned values above under the assumption that they were the sole (or first) perturbation experiment performed. In reality, of course, a line of enquiry proceeds via a series of such experiments, begging the question whether a perturbation experiment can be informative on its own but not so much if it follows another perturbation experiment. We explored this question further, by examining every possible pair of experiments (performed sequentially), and noted that there are indeed such examples. However, in the interest of continuity we do not discuss this analysis here, referring the interested reader to Supplementary S3 Table.

#### Interpreting the information gained from an experiment

We next moved beyond asking ‘how much’ information was gained from an experiment to the more subjective question of ‘what’ information was gained. To answer this, it seems natural to compare the original (wild-type) ensemble of models to the filtered ensemble that is additionally consistent with the new experiment’s results. The challenge then becomes: *how do we compare these two ensembles in a language that appeals to the biologist’s intuition?* One pragmatic approach that we devised, and illustrate here, is to identify a second experiment for which the two ensembles make markedly different predictions, and use this difference to illustrate the distinction between ensembles. For instance, consider the ‘CIC site mut.’ experiment, which we saw above to be of modest information theoretic value (Fig 3A). We also noted in Table 1 that this experiment induces a filtered ensemble with two of the eight original clusters completely ruled out and two additional clusters drastically reduced in size (from ~900 and ~700 models to 2 and 1 models respectively), suggesting that certain plausible mechanistic hypotheses were indeed ruled out by it. To interpret this further, we considered the predictions of this filtered ensemble on the ‘DL 1 site mut.’ experiment (Fig 3E) and found these to be in fair agreement with the true results from the literature [36] (Fig 3C). We then noted that the wild-type ensemble, not filtered by the ‘CIC site mut.’ experiment, is far more uncertain in its predictions about the ‘DL 1 site mut.’ experiment (Fig 3D). Thus, the ‘CIC site mut.’ experiment informs us, correctly, that mutagenizing the strongest DL site in the enhancer should not result in a significant reduction in peak *ind* levels, a point that was ambiguous in the original ensemble.

A similar approach can be adopted to interpret the information provided by other perturbation experiments. In our second example, we interpreted the ‘DL 1 site mut.’ experiment by examining the predictions of its filtered ensemble on the ‘CIC site mut.’ experiment, which according to the literature [38] shows an extension of the dorsal boundary of *ind* expression (Fig 3F) This derepression effect is much more accurately predicted by the filtered ensemble (Fig 3H), while the original ensemble’s average prediction is less definitive in predicting this effect (Fig 3G). In other words, the ‘DL 1 site mut.’ experiment informs us that CIC is an important repressor of the *ind* gene, setting up its precise dorsal boundary. For our third example, we note that the filtered ensemble of the ‘VND KO’ experiment accurately predicts that a genetic knockout of SNA will not affect the ventral boundary of *ind* expression (Fig 3I,K), while the original ensemble erroneously predicts ventral de-repression (Fig 3J). In other words, the ‘VND KO’ correctly informs us that SNA does not position the ventral boundary of *ind* expression. Thus, these three examples show how the information gained by an experiment can be interpreted by examining unique aspects of predictions of that experiment’s filtered ensemble on a second experiment.

#### Quantifying and interpreting the value of perturbation experiments on the *sim* enhancer

Similar to *ind*, single minded (*sim*) is dorso-ventral patterning gene in *D. melanogaster* that has been the subject of many biological experiments that describe the regulators of the gene, delineate its enhancer [39–41], and characterize the combinatorial action of multiple TFs and cell signaling in the formation of the precise expression pattern driven by the *sim* enhancer [42]. The *sim* gene is initially expressed at the cellular blastoderm stage in a narrow row of width equal to two cells along the dorso-ventral axis at the mesectoderm (the boundary between mesoderm and neural ectoderm) [40,41] (Fig 4A). Sim acts as a master regulator during the development of central nervous system (CNS) [43] and the confinement of its expression to the narrow line of cells is essential for the formation of the ventral midline and CNS during gastrulation [39,44]. This precise pattern of expression can be explained by a complex regulatory mechanism that involves Notch signaling [45–47]. On the ventral side, DL and Twist (TWI) activate but SNA represses the expression in the mesoderm [39,44]. Expression on the dorsal side is inhibited directly by Suppressor of hairless (Su(H)), which is the only known repressor of *sim* in the neuroectoderm [45], but is believed to have an activating influence on *sim* in the mesoderm region [45,48,49]. The sharp dorsal boundary of *sim* is formed because Notch signaling converts the ubiquitously expressed Su(H) from a repressor in dorsal regions to an activator in ventral regions exactly at the mesectoderm [37,45,48,50]. With these pieces of mechanistic information in hand, we employed GEMSTAT to model the expression driven by the *sim* enhancer. Then, we used the procedure introduced above to examine different perturbation experiments related to this enhancer reported in the literature, and quantify and interpret the ‘value’ of these experiments, after the fact. To our knowledge, this work is the first attempt to computationally model the expression of the *sim* enhancer, using the combinatorial action of TFs and signaling [45,47,51].

**Fig 4.**
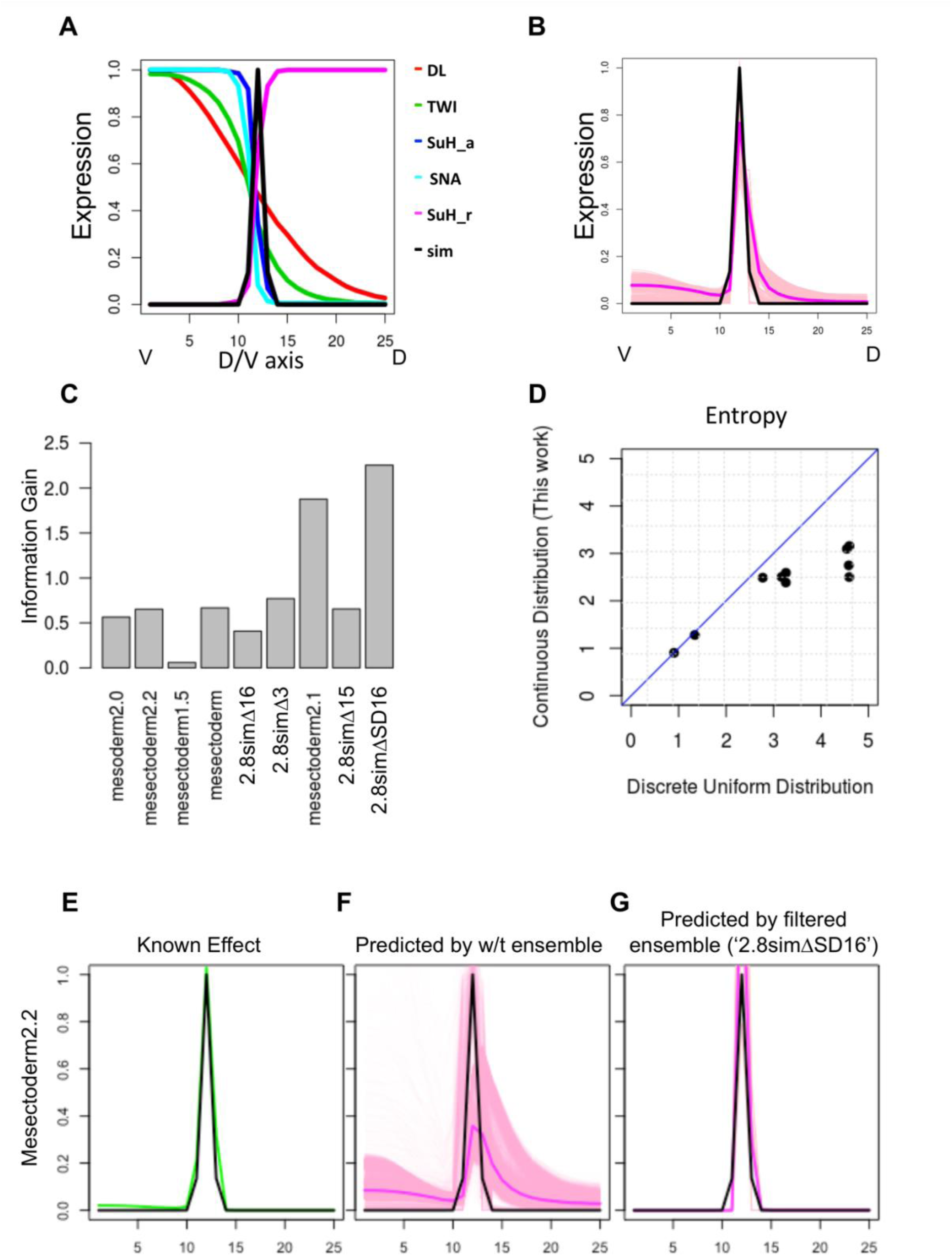
**(A)** Modeling SIM enhancer. Expression profile of sim and all TFs that are involved in sim regulation, shown for ventral-most bins 1-25 of the 50 bins along D/V axis. Su(H) is modeled as both an activator and a repressor. **(B)** Ensemble of models that predict SIM expression profile accurately. **(C)** Information Gain from different perturbation experiments. Each experiment represents the readout of a variant of the wild-type enhancer, under wild-type conditions, and is named for the variant enhancer. **(D)** Entropy of filtered ensemble for each of the nine experiments, as defined under the specially constructed probability distribution presented in this work (Y axis) or under a discrete uniform distribution (X axis). **(E-G)** The ‘Mesectoderm2.2’ variant of the *sim* enhancer [52] (S2 Table) has been observed to recapitulate the expression pattern of the wild-type enhancer (E). In contrast, majority of the models in the wild-type ensemble (F) predict expansion of the dorsal boundary, but an ensemble filtered by the ‘2.8simΔSD16’ experiment predicts the known (non-)effect correctly.

We built an ensemble of models that predicts the wild-type expression profile of *sim* accurately (Fig 4B and Methods) from its wild-type enhancer (‘2.8sim’). We then considered several experiments reported in the literature pertaining to this gene, with the goal of computing the information gain from each experiment and interpreting the information they provide. Each of the nine experiments considered is a reporter assay with a variant of the wild-type *sim* enhancer, and we used its observed readout to construct objective criteria (S2 Table) for filtering models and creating a ‘filtered ensemble’ for that experiment (S6 Fig). This allowed us to quantify the information gain score of each experiment, using the procedure described in previous sections (Fig 4C, D). This revealed that the experiment ‘2.8simΔSD16’, representing a deletion of two segments (harboring a SNA site and an E-box element respectively) from the wild-type enhancer 2.8sim, is the most informative (value 2.25), while seven of the other eight experiments are substantially less informative (about 0.5 or less). Following the procedure of the previous section, we then sought to interpret the information gained by this experiment. This was most apparent when we used the filtered ensemble of this experiment to predict the outcome of another experiment (‘mesectoderm2.2’). This second experiment is the reporter readout of a 2.2-kb sequence upstream of the early sim promoter and overlapping with the wild type enhancer 2.8sim considered above. According to the literature [52], the expression driven by this sequence is unchanged (Fig 4E) from wild-type. The filtered ensemble of the ‘2.8simΔSD16’ experiment [39] can predict this known outcome accurately (Fig 4G), while the wild-type ensemble is far more uncertain in its prediction (Fig 4F). The ‘2.8simΔSD16’ experiment tests the effect of deletion of SNA sites on the gene expression and restricting the wild-type ensemble based on results of this experiment informs us which group of SNA sites is important to set up the precise expression of the *sim* gene.

## Material and methods

### Construction of Model Ensemble

We used the GEMSTAT model from [22] with 13 different parameters, including two parameters for each of the five TFs: one pertaining to TF-DNA binding (*K_TF_*) and one to the TF’s effect on transcription rate (*α_TF_*). Moreover, the model has one parameter for DL-ZLD cooperativity (*w_DL_*_–*Zld*_), and another parameter reflecting the baseline transcriptional rate (*q_BTM_*). The repressor CIC has a uniform dorso-ventral expression profile but its repressive effect is attenuated in the neuroectodermal region by locally activated ERK, through a reduction in CIC-DNA binding; this effect, as modeled in [21], is represented by a free parameter (*Cic_att_*). The expression pattern of each TF and the *ind* gene was scaled in the range of zero to one. Each expression profile is represented by a 50-dimensional vector (S1A Fig), with dimensions corresponding to equally spaced bins along the D/V axis (bin 1 = ventral end). The *ind* gene is expressed in only 5 to 7 of these bins (bins 22-28 with the peak of expression at bin 25).

To construct the ensemble, we sampled models from the 13-dimensional parameter space following the procedure of Samee et al. [21]. We divided the range of each parameter into two halves (using log scale for *K* parameters of all TFs, *α* parameters of repressors and for *q_BTM_*, and scale fo: *α* of all activators, *w_DL_*_–*Zld*_ and *Cic_att_*), and sampled 1000 points in each cell of the 13-dimensional space, for a total of ~8 million models. Such a dense sampling was possible because the number of parameters (13) is modest. Among these randomly generated models, we retained those with SSE (sum of squared errors) score between the real and predicted *ind* profile less than 10%. This higher initial threshold allows us to get good initial points. We then used GEMSTAT’s optimization routine to locally optimize the initial ensemble, following which we filtered for models that have SSE score less than 5%. (This stricter error threshold was determined by a visual inspection of many examples, as one that allows the main spatial pattern of expression to be preserved in the predictions meeting that threshold.) We call the resulting collection the wild-type ensemble of models. It contains more than 5000 distinct parameter settings from about 600 different regions, and all of these models make good predictions on the wild-type data, as per visual inspection (Fig S1B).

### Construction of Probability Distribution over Ensemble of Models

Each model is represented by its parameter vector, and we first scaled the parameters using min-max scaling to place all the dimensions on the same scale of 0-1. The models were then clustered based on a multivariate density estimation method, Mclust [53,54], using its R implementation. This method approximates the complete collection of models as a (generally non-uniform) mixture of Gaussian distributions, each component Gaussian representing a cluster, with its own mean (cluster center) and covariance matrix (S5 Fig), while simultaneously determining the optimal number of clusters. Next, we modeled each cluster *C* as a uniform mixture of *n_C_*=|*C*| Gaussian distributions with each of the *n_C_* models of the cluster as mean, and a common covariance matrix Σ_*C*_ estimated for the cluster by the Mclust method. The following formula describes the probability distribution over the space of models *θ*:

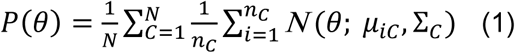

where *N* is the number of clusters, *C* indexes these clusters, *n_C_* is the number of models in cluster *C*, *μ_iC_* is the *i*^th^ model in the cluster *C*, and *Σ_C_* is the covariance matrix of cluster *C*.

Since models in the ensemble are not simply random samples of the probability distribution, but a collection of local optima obtained from initial random samples as seeds, we did not use a standard density estimation technique to build a density function. (See Discussion for our choice of the above methodology as opposed to more sophisticated sampling techniques.) We expected the distribution to reflect the fact that each model is a local optimum in the landscape of models scored by SSE measure and they group into several clusters. These clusters reflect different hypotheses for the biological mechanism of the underlying system and we desired that the probability density function put equal weights on them. Within each cluster, it is possible to observe several equally good local optima (peaks of the SSE landscape); we selected each such optimum as a local peak for the probability density function as well, constructing a Gaussian Mixture with a fixed covariance matrix to represent a cluster, with component Gaussians centered on the optimized models in that cluster.

### Entropy of Probability Distribution over Ensemble

We calculated the entropy function using a discrete version of the probability density function in (1). The discrete probability *p_iC_* of the model *i* in the cluster *C* is set to be proportional to the value of the continuous probability density function at the location of the model. Each *p_iC_* receives contributions from all Gaussians in the mixture model. We set the constant of proportionality such that for each cluster *C* we have 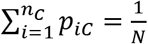 where *N* is the number of clusters. We then estimated the entropy of this discrete probability density function using the formula 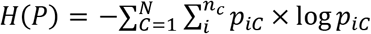 [55].

### Information Gain from an Experiment

Suppose we are given an ensemble of models obtained from a set of experiments and have calculated its probability distribution. We are then given the results of a new experiment. This is typically in the form of gene expression level(s) driven by an enhancer sequence in one or more cellular contexts described by their TF concentration profiles. We assess if the predictions made by any model in the ensemble are consistent with these results, by comparing the model’s predictions to the observed gene expression levels through a single goodness-of-fit score such as SSE (sum of squared error), and discard that model if this score is worse than a pre-determined threshold. Repeating this process for each model in the ensemble, we obtain a ‘filtered ensemble’ that is a subset of the original ensemble. We then modify the probability distribution over the filtered ensemble by (a) retaining the cluster assignments and cluster covariance matrices of the original ensemble, but (b) removing any cluster that has no remaining models and readjusting weights of remaining clusters to add to 1, and (c) redefining the Gaussian Mixture Model for each cluster to have component Gaussian distributions centered at each model remaining in that cluster (also with uniform weights). Having thus defined a probability distribution over the filtered ensemble, we calculate its entropy as above and assign the difference of entropy between the (distributions over the) original and filtered ensemble as the information theoretic ‘value’ of the new experiment.

### Data sets for modeling *sim* enhancer

We constructed a one-dimensional vector to describe the expression readout of the *sim* enhancer along the D/V axis, as recorded in the literature. This vector (and other expression vectors described here) has 25 dimensions, representing equally spaced positions (‘bins’) along the axis from the ventral end to the mid-point of the axis. (These are the same as bins 1-25 of the 50 bins considered in previous sections.) The expression in each bin has a value between 0 and 1 that corresponds to the relative amount of gene expression observed in that bin. The expression profile of the wild-type *sim* enhancer was represented as a Gaussian curve that has its peak at the location where SNA expression changes from high to low (12^th^ bin from ventral end), i.e., at the known location of the *sim* expression peak. The variance of the Gaussian is set to be small enough that the ‘width’ of the expression profile is similar to the narrow domain in which SNA goes from high to low (Fig 4A). We obtained TF (protein) expression profiles of DL, TWI, and SNA from Zinzen et al. [17] and represented them in the same 25-dimensional vector format as above (Fig 4A). We considered DL and TWI as activators and SNA as a repressor. The other important regulatory input considered was Su(H), which is a maternal protein uniformly expressed across the D/V axis. It is believed to be a repressor, but Notch signaling activated by the effect of SNA on Notch-Delta endocytosis switches the role of Su(H) from a repressor to an activator [45–47] in domains of SNA expression (mesoderm). Since GEMSTAT does not allow for such a ‘role-switch’ for any TF, we separated the uniform expression profile of Su(H) into two separate profiles (vectors), one for each role: an ‘activator Su(H)’ with an expression profile similar to SNA but extended to include the mesectodermal positions and a ‘repressor Su(H)’ with its complementary profile. In this manner we capture the prior knowledge of the ‘role-switch’ of Su(H) at the peak expression of *sim*. The *sim* enhancer sequence and TF motifs, required by GEMSTAT, were taken from Fly Factor Survey [56].

## Discussion

Determining regulatory mechanisms shaping the spatio-temporal pattern of a gene of interest is a tedious process. While high throughput technologies provide helpful clues and narrow the space of possibilities, the ‘gold standards’ for demonstrating the regulatory influence of a transcription factor on a gene – a combination of TF knockout or overexpression (and observed effects on gene expression), TF-DNA binding assays, site mutagenesis and rescue experiments – involve substantial investments. Guidance about the most insightful experiments to perform, given current knowledge about the gene’s regulation, can thus be highly beneficial. Typically, such choices are made by the biologist by relying on their intuition. We asked ourselves if the process of designing experiments to gain deeper understanding of a gene’s regulatory mechanisms may be made systematic. This immediately presented two major conceptual challenge: first, how do we formalize what is ‘current knowledge’ about the gene’s regulation, and second, how do we measure how insightful or informative an experiment is? Answers to these questions appear to be necessary before we could systematize the process of experiment selection or design, mentioned above. In this manuscript, we take present a possible solution to these challenging problems by making use of a previously established quantitative modeling framework that relates trans- and cis-regulatory information to gene expression levels, and combining the framework with ensemble modeling and information theoretic ideas. In the future, our approach can be combined with well-established ideas in statistical experiment design [28,29] to develop a full-fledged formal approach to investigation of gene regulatory mechanisms.

We approached the goal of formalizing current knowledge about a gene’s regulation by using the GEMSTAT framework of gene expression modeling. (Other related models, e.g., [13], would also have been similarly usable.) Here, current knowledge of a gene’s enhancer sequence(s) and its known regulators (TFs) is encoded into a mathematical function that is consistent with data on the gene’s and TFs’ expression levels in multiple conditions or cell types. This not only forces the qualitative knowledge of regulators into a precise quantitative form, it also explicitly captures complexities and subtleties associated with combinatorial action of multiple TFs. Furthermore, the model has free parameters representing important but often uncharacterized biochemical properties of the regulators, viz., free energy of DNA binding and strength of regulatory influence, and the modeling step involves assigning values to these parameters so as to match available data. In this step, one is often faced with many distinct parameter settings that appear equally plausible in light of available data, and these different parameterizations represent ambiguities in current mechanistic understanding of the gene’s regulation, even when the likely regulators are qualitatively characterized. In our approach, such ambiguities are explicitly catalogued in the form of an ensemble of models consistent with data.

To address the other conceptual challenge mentioned above (‘how informative is an experiment?’), we compared the ensemble representing prior knowledge/data to that representing new experimental data in addition to the prior information. It was natural to consider using the information theoretic ideas of entropy and information gain for purposes of this comparison. We therefore devised an approach to define a probability distribution over models in the ensemble, and to estimate the entropy of the distribution; the information gain was then defined as the difference in entropy of the two ensembles.

We demonstrated the use of our approach in the context of two genes in early fruitfly development – *ind* and *sim* – whose regulatory mechanisms have been studied through several perturbation experiments (TF knockouts, site mutagenesis, variant enhancers, etc.) reported in the literature. In each case, we started with the wild-type enhancer and likely regulators as ‘current knowledge’ and (retroactively) quantified how informative each of the perturbation experiments is. We also presented objective observations about each experiment that suggest the specific insights it added to our understanding of the gene’s regulatory mechanisms. In the case of *ind*, we additionally applied our experiment-scoring framework in a more controlled, semi-synthetic setting, where real data on the gene were used to first select a unique model as the underlying ‘truth’, and used to provide the results of *in silico* perturbation experiments. We note however that the information gain values computed by our method are not comparable across different studies, e.g., between the *ind* and *sim* studies considered here; they are only comparable across different experiments for the same gene, when evaluating those experiments for additional insights over a common set of current data/knowledge.

Methodologically, an important feature of our approach was the generation of ensemble by uniform sampling in the multi-dimensional space, followed by optimization, as was done in [21]. Alternative sampling algorithms can be used to generate a large sample from optimal regions of the parameter space. Sampling methods such as Bayesian optimization techniques can be used to efficiently search for optimal parameters of any given model with nonlinear cost functions [57]. However, in practice these sampled optimal solutions may not be representative of every possible region of the parameter space with similar goodness of fit. One example of such a Bayesian optimization algorithm is the spearmint package. Spearmint, when applied to our problem, produced only a few optimal models (data not shown). Building a large ensemble of models that represents every locally optimal region requires running the method multiple times, which is very slow, due to slow convergence time [58]. The spearmint method yields only few data points as the result of optimization and they are usually close to each other. Unless we perform a detailed sampling around those points or combine such techniques with a more global sampling approach, we do not have a diverse ensemble to work with. On the other hand, since the number of parameters in our model is small (less than 20), we could afford to do a dense uniform sampling of the parameter space with our approach.

We believe that the framework established in this work can be used in future work to formalize experiment design strategies for gene regulation studies. For instance, we may compute the expected information gain [59] of various possible future experiments and select the best one. Given a candidate future experiment, we may first use each in the current ensemble to predict its outcome, compute the resulting information gain, and then compute an expectation of this value over the entire ensemble, using the probability distribution introduced in our work.

## Supporting information

**S_File.pdf: Supplementary Figures and Tables**

